# Vocabulary recurrence, rather than biological grounding, explains tokenization transfer in protein function prediction

**DOI:** 10.64898/2026.11.16.767127

**Authors:** Tolulope S. Adeyina, Jonathon E. Mohl, Jason McDermott, Ali Aslam, Kenneth Onyedibe, Muhammed Idris, Mustapha Olayemi Oloko-Oba

**Affiliations:** Morehouse School of Medicine, Atlanta, GA, USA; The University of Texas at El Paso, El Paso, TX, USA; Pacific Northwest National Laboratory

**Keywords:** protein function prediction, tokenization, Gene Ontology, data leakage, tryptic peptides, evaluation methodology, out-of-vocabulary

## Abstract

Sequence-based protein function prediction requires a choice of tokenization. K-mers, the common default, correspond to no biological unit; enzymatic digestion offers a principled alternative, since tryptic peptides are the fragments proteases generate and mass spectrometry measures. We asked whether this grounding improves transfer to unseen proteins. Using six Gene Ontology terms and 23,723 UniProt proteins, we compared tryptic peptides, peptide topics (groups of related peptides), 3-mers, and amino-acid composition, with ESM-2 embeddings and homology transfer as reference points. Peptide performance followed vocabulary coverage: 92.2% of peptide tokens in test proteins were never seen in training, whereas almost every 3-mer had been seen. As a result, 38.9% of test proteins contained no peptide the model had seen, and with nothing to work from, the classifier gave all of them the same answer. The remaining 61.1% of proteins were classified almost perfectly with a random forest; however, because of this uncovered fraction, tryptic peptides scored lower overall than 3-mers under all three classifiers tested (macro-F1 0.640 against 0.690). These results suggest that vocabulary recurrence matters more than biological grounding for tokenization transfer.

## 1. Introduction

The gap between sequenced and characterized proteins continues to widen: UniProtKB held approximately 246 million sequence records at release 2024_04 (1), of which only about 570,000 were manually reviewed in Swiss-Prot. Computational annotation also carries real risk: across 37 enzyme families from six superfamilies, misannotation of molecular function averaged 5 to 63% per superfamily in GenBank NR, TrEMBL and KEGG and exceeded 80% in at least one of these databases for 10 of the families, while remaining close to zero for most families in curated Swiss-Prot (2). Assigning function from sequence is therefore still an open problem. The Gene Ontology (3,4) provides the vocabulary in which predictions are expressed, and the Critical Assessment of Functional Annotation (CAFA) set the benchmark framework across its first three rounds, CAFA1 through CAFA3 (5–7), with further rounds following since.

Most sequence-based methods that require tokenization begin by breaking the protein into smaller units, or tokens, from which a representation is learned. That choice is usually made by convention rather than by argument. The prevailing default is the fixed-length *k*-mer, every overlapping window of *k* residues, embedded using methods borrowed from natural language processing. ProtVec, the canonical example, learns a numerical vector (an embedding) for each overlapping 3-mer using the skip-gram method, in which a model is trained to predict the tokens that appear near a given token in the sequence, so that tokens found in similar contexts receive similar vectors. It reaches an average family classification accuracy of 93% across 324,018 Swiss-Prot sequences in 7,027 families (8). *K*-mers are simple, exhaustive and bounded, yet biologically arbitrary: a three-residue window corresponds to no structural element, no functional motif, and no entity handled by cellular machinery or analytical instruments. It is an artefact of the encoding rather than a fact about the protein.

Enzymatic digestion suggests a more principled alternative. Trypsin cleaves exclusively C-terminal to lysine and arginine residues (9), conventionally excepting sites followed by proline, a rule whose empirical status has itself been re-examined at scale (10). The resulting peptides are chemically real, since they are what proteolysis produces. They are also experimentally real: in bottom-up proteomics, proteins are digested, usually with trypsin, and it is the peptides rather than the intact proteins that are separated, fragmented and sequenced by mass spectrometry (11,12). A representation built on tryptic peptides would be grounded twice over, biochemically in an actual reaction and experimentally in a measurable analyte. In principle it would let function predictions be anchored to peptides observable in a sample rather than to windows that exist only in software.

Variable-length tokenization of proteins is not new. Subword schemes such as peptide-pair encoding (13), ProtGPT2’s byte-pair vocabulary (14), and evoBPE (15) use probabilistic or corpus-frequency merges to construct variable-length tokens. Systematic benchmarks indicate that small subword vocabularies (50–200 elements) perform best while larger vocabularies degrade accuracy (16), and that statistical subwords align poorly with functional domain boundaries (17). Crucially, these approaches derive units statistically from corpus frequency rather than biochemistry. While biologically defined units such as protein domains have been embedded with Word2Vec (18), tokens defined by enzymatic reactions remain untested. Here, we introduce tokens defined by an enzymatic reaction, tryptic peptides, as a representation for protein function prediction, and test how well they transfer to unseen proteins.

There is a reason such an evaluation needs care. For any tokenization to support prediction on unseen proteins, its vocabulary must recur : tokens learned in training have to reappear in proteins never seen. K-mers satisfy this trivially, since the token space is bounded and any substantial corpus samples it densely, so the open-vocabulary problem that motivated subword encoding in language (19) does not arise. Enzymatic peptides carry no such guarantee. Their length is set by the spacing of cleavage sites, and a single substitution inside the peptide or at either boundary changes their identity entirely. Existing work points the same way: byte-pair encoding is valued for proteins because it reduces vocabulary sparsity and handles unseen tokens by construction, fixed-length k-mer tokenization is criticised for severe sparsity at large k (20), and tokenization choice measurably affects transformer performance on biological sequences (21). What has not been measured is how often an enzymatically defined vocabulary recurs in unseen proteins, or what happens to a classifier when it does not.

Because exact tryptic peptides may recur too rarely to support prediction, we also include a coarser representation that groups similar peptides together, which we call peptide topics. Individual peptides are embedded with skip-gram Word2Vec and clustered into groups, with each protein represented by the distribution of its peptides across these topics rather than by exact peptide identities. This serves two purposes. It tests whether pooling related peptides can recover the recurrence that exact peptides lack, since different peptides in the same topic are treated as the same feature. It also asks whether the groups correspond to functional units, which would restore some of the biological meaning that motivated enzymatic tokenization. Peptides absent from the training vocabulary cannot be placed in a topic, so out-of-vocabulary peptides are tallied in a dedicated count column.

Evaluating any of these representations on unseen proteins also requires controlling for sequence redundancy, now recognised as a primary source of inflated performance estimates in biological machine learning (22–24). Reported accuracy can fall substantially once redundancy is removed, in one illustration for protein–protein interface prediction from 0.92 to 0.72 AUC (24), and homology-aware partitioning tools have been built to address it (25,26). CAFA stratifies its benchmarks by maximum sequence identity to the training set because difficulty depends on it (6), and homology transfer by sequence search remains a baseline that learned methods struggle to beat (7,27).

We therefore asked whether a biologically grounded tokenization transfers better to unseen proteins than an arbitrary one. We evaluate four representations, tryptic peptides, peptide topics, 3-mers and amino-acid composition, for Gene Ontology term prediction across six terms and 23,723 UniProt proteins, with two reference points: mean-pooled ESM-2 embeddings (28), which bound what is recoverable from sequence and are competitive with dedicated function predictors (29,30), and homology transfer by sequence search (31). Our contributions are as follows:

1. We benchmark against two independent reference points, ESM-2 and homology transfer, confirming that functional signal is recoverable across the split, and show that tokenization performance trends hold across three distinct classifiers.
2. We measure vocabulary recurrence for enzymatic peptides against 3-mers, showing that extreme out-of-vocabulary rates are intrinsic to enzymatic cleavage and not an artefact of frequency pruning.
3. We separate test proteins by vocabulary coverage, demonstrating that peptide classification reduces to exact matching on covered proteins and uniform default predictions on uncovered ones.
4. We evaluate what peptide topics capture, checking topic purity against GO class and revealing that frequent topics reflect database composition and sequence fragments rather than functional motifs.
5. We evaluate cluster-disjoint splitting, showing that standard greedy clustering leaves substantial pairwise identity across partitions, and provide an empirical audit against measured identity.

## 2. Materials and Methods

### 2.1 Dataset construction

Protein sequences were retrieved from UniProtKB through the REST streaming endpoint using unrestricted Gene Ontology term queries of the form (go:XXXXXXX). No reviewed-status filter and no free-text filter were applied, so both Swiss-Prot and TrEMBL entries were returned. Six GO terms were used: GO:0002933 (lipid hydroxylation, biological process), GO:0004608 (phosphatidylethanolamine N-methyltransferase activity, molecular function), GO:0004653 (polypeptide N-acetylgalactosaminyltransferase activity, molecular function), GO:0004713 (protein tyrosine kinase activity, molecular function), GO:0015977 (carbon fixation, biological process) and GO:0043885 (anaerobic carbon-monoxide dehydrogenase activity, molecular function).

Retrieval was performed on 15 August 2026 against UniProt release 2026_02, returning 331, 3,409, 26,446, 269,123, 58,728 and 4,985 sequences for the six terms respectively. Downloads were validated against the record count reported by the UniProt search API, and any file deviating by more than 1% was retrieved again, falling back to cursor pagination where the streaming endpoint terminated early.

Sequences were parsed from FASTA headers matching both sp| and tr| prefixes, capturing accession, entry name, description, organism and fragment status. Sequences shorter than 30 or longer than 5,000 residues were excluded, as were sequences containing characters outside the standard and ambiguity amino-acid alphabet.

Each term was then capped at 5,000 sequences by random sampling with a fixed seed. Two of the six terms overlap biologically, since anaerobic carbon-monoxide dehydrogenase performs carbon fixation through the Wood–Ljungdahl pathway. Because an identical input sequence cannot have two correct single-label answers, accessions appearing under more than one term were removed rather than assigned arbitrarily. One accession met this criterion after sampling (P27989, *Neomoorella thermoacetica* carbon monoxide dehydrogenase/acetyl-CoA synthase subunit beta), one protein of 23,723, reducing those two classes by one protein each. This gave a corpus of 23,723 proteins: GO:0002933 (331), GO:0004608 (3,409), GO:0004653 (5,000), GO:0004713 (5,000), GO:0015977 (4,999) and GO:0043885 (4,984). Fragmentary entries made up 15.0% of the corpus and were kept, since excluding them would bias the corpus toward well-studied proteins; their distribution across classes is uneven, from 1.2% of GO:0043885 to 25.3% of GO:0015977. The corpus is overwhelmingly unreviewed, with 23,410 TrEMBL entries against 313 Swiss-Prot (98.7%), so annotations should be understood as predominantly electronically inferred.

The six terms were chosen as a targeted case study rather than a representative sample of GO. They differ widely in size (331 to 269,123 sequences returned) and in kind, with four molecular-function terms and two biological-process terms, so we treat them as a case study and not as a representative sample of GO. The two process terms behave differently: GO:0002933 is dominated by a single homology component (Table 1) and so behaves like a family, whereas GO:0015977 spans unrelated folds. Its members include ribulose-bisphosphate carboxylase large and small subunits, phosphoenolpyruvate carboxylase, propionyl-CoA carboxylase, phosphoribulokinase, fructose-1,6-bisphosphate aldolase and glyceraldehyde-3-phosphate dehydrogenase, which are unrelated folds sharing a metabolic role. We therefore do not describe the label set as six disjoint superfamilies.

**Table 1.** Homology component structure by class.

| **GO term** | **Components** | **Proteins** | **Largest component share** |
|---|---|---|---|
| GO:0002933 | 9 | 331 | 0.882 |
| GO:0004608 | 180 | 3,409 | 0.420 |
| GO:0004653 | 440 | 5,000 | 0.871 |
| GO:0004713 | 1,025 | 5,000 | 0.231 |
| GO:0015977 | 211 | 4,999 | 0.321 |
| GO:0043885 | 90 | 4,984 | 0.288 |

### 2.2 Homology structure and data splitting

All 23,723 sequences were searched against each other with MMseqs2 (31),(MMseq2 is a fast sequence search and clustering tool for protein and nucleotide sequences), using the easy-search module at sensitivity 7.5, E-value threshold 1e^−3^, retaining up to 1,000 hits per query and requiring 80% bidirectional coverage.

We first attempted a split with a structural guarantee of homology separation. An undirected graph was built over all sequences, with an edge between any pair at 30% identity or above and 80% bidirectional coverage or above, and connected components taken as grouping units. Components are transitively closed: if A is related to B and B to C, all three sit in one group and necessarily fall on the same side of any group-aware partition.

This produced 1,954 components, with the largest holding 18.4% of the corpus. The components are distributed unevenly across classes (Table 1). Three classes are dominated by a single component. Because a component is atomic under any group-aware partitioner, such a class is assigned almost wholly to training or test, precluding stratified allocation. We evaluated all five folds of a stratified group partition, requiring every class to retain at least 30 test proteins and no class to deviate by more than 20 percentage points from the target test fraction. None qualified: three folds left a class with no test proteins at all, and the remaining two placed GO:0002933 at 6 test proteins and at 6 training proteins, respectively. This limitation is intrinsic to the corpus: where functional classes collapse into single homology components, homology-disjoint splitting and class balance are mutually unsatisfiable.

We therefore fell back to the standard approach. MMseqs2 easy-cluster at 30% minimum sequence identity and 80% bidirectional coverage produced 2,636 clusters, a 9.0-fold redundancy, used as grouping units in a five-fold stratified group partition. Applying the same two criteria, exactly one of the five folds qualified. Selection was therefore forced rather than chosen: of ten candidate partitions across both grouping schemes, one satisfies the criteria. This gave 20,542 training and 3,181 test proteins with no cluster shared between partitions, asserted programmatically. Class support is uneven by construction (Table 2), and per-class results are reported throughout.

**Table 2.** Split composition.

| **GO term** | **Train** | **Test** | **Test fraction** |
|---|---|---|---|
| GO:0002933 | 296 | 35 | 0.106 |
| GO:0004608 | 3,278 | 131 | 0.038 |
| GO:0004653 | 4,710 | 290 | 0.058 |
| GO:0004713 | 4,144 | 856 | 0.171 |
| GO:0015977 | 4,631 | 368 | 0.074 |
| GO:0043885 | 3,483 | 1,501 | 0.301 |

Because cluster-disjointness does not by itself guarantee low pairwise identity between partitions (Section 4.3), we measured it directly. For every test protein we computed the maximum fractional identity to any training protein over alignments covering at least 80% of both sequences, and we report performance against this measured identity alongside aggregate performance (Sections 3.4 and 3.5). We therefore do not describe this split as homology-disjoint.

### 2.3 Tokenization and representations

Sequences were digested in silico at positions C-terminal to lysine or arginine, excluding sites immediately followed by proline (9,10). No missed cleavages were allowed in the primary analysis. Peptides shorter than 6 or longer than 40 residues were discarded. All overlapping 3-mers were extracted for the k-mer arm, giving a bounded vocabulary; 8,792 distinct 3-mers were observed (the excess over 20³ arising from the ambiguity codes retained above).

All unsupervised representations were fitted exclusively on training sequences; test sequences were never seen during vocabulary construction, embedding, or clustering. Amino-acid composition served as the baseline floor: relative frequencies of the 20 standard amino acids (20 features), omitting sequence order and learned tokens. Peptide and 3-mer embeddings were trained using skip-gram Word2Vec (100 dimensions, window size 5, 20 epochs, minimum token count 5) fitted on the training partition with a single worker and a fixed seed to guarantee determinism. Each protein was represented as the mean embedding of its in-vocabulary tokens; proteins lacking training-vocabulary tokens received an all-zero vector, an outcome we quantify directly rather than mask.

For the topic arm, we clustered the 100-dimensional training Word2Vec vectors of in-vocabulary peptides using MiniBatchKMeans. The number of clusters was set to the square root of the training vocabulary size (*k* = 124). Each protein was then represented by a histogram counting how many of its constituent peptides fell into each of the 124 clusters. One additional feature recorded the count of out-of-vocabulary peptides, ensuring every protein received a non-zero representation across 125 features.

The protein language model arm used mean-pooled residue embeddings from ESM-2 650M (28), computed in half precision with beginning- and end-of-sequence tokens excluded from pooling, yielding 1,280 features. Sequences were truncated at 1,022 residues, affecting 1,393 of 23,723 proteins (5.9%).

Truncation was distributed unevenly, affecting 20.5% of GO:0004713 (median length 705 residues) compared to 6.3% of GO:0004608, 1.7% of GO:0015977, 1.2% of GO:0004653, 0.2% of GO:0043885, and no GO:0002933 proteins. Because ESM-2 was pretrained on UniRef (which contains these sequences), this arm serves as an optimistic reference bound to demonstrate that the functional signal is recoverable from sequence.

For the homology-transfer baseline (5,27), test sequences were searched against the training partition using MMseqs2 easy-search (sensitivity 7.5, E-value threshold 10^5^, up to 300 hits per query). Each test protein was assigned the GO term of its highest-scoring training hit by bit score. The permissive E-value threshold (10^5^) was chosen deliberately so that the identity distribution of best hits could be fully characterized without artificial truncation.

### 2.4 Classification and evaluation

All representations were classified with a random forest of 500 trees using balanced subsample class weighting, held fixed across arms so that differences reflect the representation rather than the model, and scored on the identical frozen test partition. To check that our conclusions do not depend on this choice, we repeated the comparison with logistic regression (standardized features, balanced class weights) and histogram-based gradient-boosted trees (200 iterations, balanced sample weights), using default settings for all remaining parameters and no tuning.

Evaluated metrics include accuracy, balanced accuracy, macro-averaged precision, recall, macro-F1, macro-averaged area under the precision-recall curve (macro-AUPR), and one-versus-rest macro ROC-AUC. We emphasize macro-F1 alongside balanced accuracy and macro-AUPR because class imbalance can penalize macro-F1 independently of representation quality (Section 3.1). All macro-averages are computed over the full set of six class labels rather than only the classes predicted by a given model. We also report classification accuracy conditioned on per-protein vocabulary coverage.

### 2.5 Implementation and availability

The pipeline is implemented as a single Jupyter notebook running end to end from data retrieval to final tables, with per-stage checkpointing. All random seeds are fixed at 42. Software versions were Python 3.12.13, numpy 2.0.2, pandas 2.2.2, scipy 1.16.3, scikit-learn 1.6.1, gensim 4.4.0, transformers 5.13.1, torch 2.11.0+cu128, and MMseqs2 build 17b688d21dda57fc5f5b7286ecba7ec003d4717f. Computation used a Google Colab instance with an NVIDIA Tesla T4 GPU. Code and derived result tables are available at https://github.com/adeyinatolulope/protein_function_model_OOV. Sequence data are not redistributed and can be retrieved from UniProt release 2026_02 using the queries in Section 2.1.

## 3. Results

### 3.1 The signal is recoverable across this split

Before attributing any difference between tokenizations to the tokenization itself, we first tested whether these six classes can be separated on this split at all. If the functional signal is recoverable from sequence, methods that do not rely on a learned tokenization should perform well, and any shortfall among the learned tokenizations can be traced to the representation rather than the task.

We evaluated two reference methods on the frozen test set (n=3,181) alongside the four representations and two trivial baselines. Homology transfer assigns each test protein the label of its highest-scoring training hit by sequence search. For ESM-2, we mean-pooled residue embeddings from the 650M-parameter protein language model across each protein (1,280 features) and trained the same random forest classifier used for the other arms (Table 3).

**Table 3.** Classification performance on the frozen test set (n = 3,181).

| **Representation** | **Features** | **Accuracy** | **Balanced acc.** | **Macro-F1** | **Macro-precision** | **Macro-recall** | **Macro-AUPR** | **ROC-AUC** |
|---|---|---|---|---|---|---|---|---|
| Homology transfer | – | 0.9965 | 0.9940 | 0.9946 | – | – | – | – |
| ESM-2 650M | 1,280 | 0.9717 | 0.8237 | 0.8280 | 0.9737 | 0.8237 | 0.9967 | 0.9995 |
| 3-mer Word2Vec | 100 | 0.8362 | 0.6771 | 0.6896 | 0.8480 | 0.6771 | 0.7135 | 0.9054 |
| Tryptic peptide Word2Vec | 100 | 0.7265 | 0.6048 | 0.6396 | 0.9130 | 0.6048 | 0.6501 | 0.9087 |
| Peptide topics | 125 | 0.6445 | 0.6011 | 0.5740 | 0.6017 | 0.6011 | 0.6526 | 0.8660 |
| Amino-acid composition | 20 | 0.5876 | 0.5870 | 0.5614 | 0.7537 | 0.5870 | 0.6494 | 0.8764 |
| Stratified baseline | – | 0.1883 | 0.1666 | 0.1388 | – | – | – | – |
| Most-frequent baseline | – | 0.0912 | 0.1667 | 0.0278 | – | – | – | – |

Both reference methods confirmed that the signal needed to separate these six classes is present in sequence and recoverable across this split. Homology transfer reached a macro-F1 of 0.995, and ESM-2 reached an accuracy of 0.972 and a macro-AUPR of 0.997 (macro-F1 0.828). Both exceeded every learned tokenization, the strongest of which, overlapping 3-mers, reached a macro-F1 of 0.690.

One small class lowers macro-F1 across all feature-based methods. GO:0002933 has only 35 test proteins, and the peptide, 3-mer, amino-acid composition, and ESM-2 arms each recover only 2 of them (recall 0.057). For ESM-2, this reflects the random forest decision threshold rather than the representation: its precision on GO:0002933 is 1.000, the mean probability assigned to the true class is 0.287 (ranking the true class second of six by median), and replacing the random forest with logistic regression raises ESM-2 macro-F1 to 0.987. We therefore report balanced accuracy and macro-AUPR alongside macro-F1 throughout.

Any shortfall of a learned tokenization therefore stems from the tokenization itself rather than task difficulty. Moreover, the aggregate gap between 3-mers and tryptic peptides (macro-F1 0.690 vs. 0.640) masks a sharp bimodal split in how peptide predictions are generated.

### 3.2 Peptide performance tracks vocabulary coverage

To understand the difference between the peptide and 3-mer arms, we examined how often tokens recur across proteins. If an enzymatic vocabulary rarely recurs, most peptides in a held-out protein will be absent from the training vocabulary, and classification accuracy should depend on whether a protein contains any known peptide at all rather than on how well function is inferred from its sequence.

We measured the fraction of peptide tokens in test proteins that were absent from the training vocabulary, compared this against the 3-mer arm (Figure 1), and partitioned test proteins by whether they contained at least one training-vocabulary peptide (Table 4). We also refitted the peptide arm across three minimum-count thresholds to verify that our findings do not depend on frequency pruning (Table 5).

**Figure 1.**
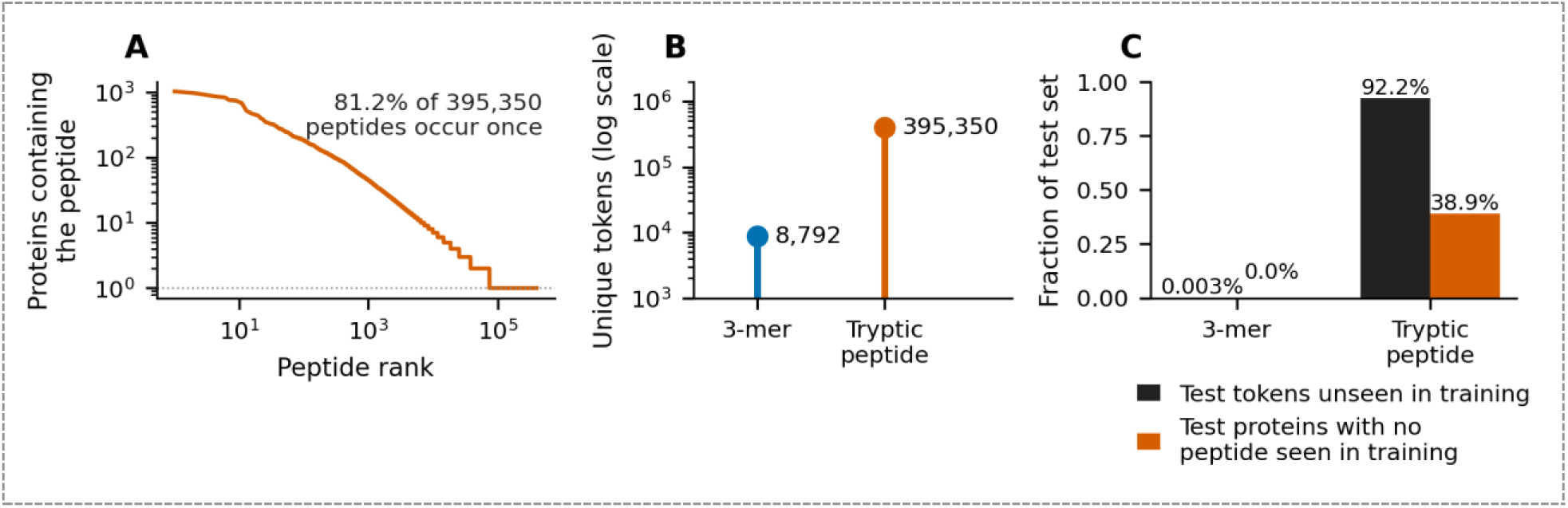
The tryptic peptide vocabulary does not recur. (A) Number of proteins containing each tryptic peptide, ranked by frequency; most of the 395,350 peptides occur in a single protein. (B) The peptide vocabulary is about 45-fold larger than the 3-mer vocabulary (395,350 against 8,792 unique tokens; log scale). (C) In the test set, 92.2% of peptide tokens were not seen in training, against 0.003% of 3-mer tokens, and 38.9% of test proteins contained no peptide seen in training, against none for 3-mers.

**Table 4.** Peptide-arm accuracy by class, split by vocabulary coverage.

| **Class** | **n** | **n zero-vector** | **Accuracy (covered)** | **Accuracy (zero-vector)** |
|---|---|---|---|---|
| GO:0002933 | 35 | 33 | 1.000 | 0.0 |
| GO:0004608 | 131 | 36 | 1.000 | 0.0 |
| GO:0004653 | 290 | 51 | 1.000 | 0.0 |
| GO:0004713 | 856 | 370 | 1.000 | 1.0 |
| GO:0015977 | 368 | 230 | 0.978 | 0.0 |
| GO:0043885 | 1,501 | 516 | 0.999 | 0.0 |

**Table 5.** Effect of the Word2Vec minimum-count threshold on peptide coverage and performance.

| **min_count** | **Vocabulary** | **Token OOV** | **Zero-vector proteins** | **Macro-F1** |
|---|---|---|---|---|
| 1 | 331,365 | 0.831 | 663 (20.8%) | 0.5593 |
| 2 | 61,478 | 0.886 | 881 (27.7%) | 0.6648 |
| 5 | 15,304 | 0.922 | 1,236 (38.9%) | 0.6396 |

Across the corpus, in silico tryptic digestion produced 395,350 distinct peptides, of which 81.2% occurred in only a single protein, leaving 15,304 peptides in the training vocabulary after applying the minimum-count threshold of 5 (Figure 1A, B). On held-out test proteins, 92.2% of peptide tokens were absent from the training vocabulary, compared to just 0.003% for the 3-mer arm, whose vocabulary is bounded at 8,792 observed tokens (Figure 1C). Consequently, 1,236 of the 3,181 test proteins (38.9%) contained no training-vocabulary peptide at all, compared to 0.0% for 3-mers.

Peptide-arm accuracy followed this vocabulary coverage in an all-or-nothing pattern. Because the 1,236 uncovered test proteins contained no known tokens, each received an identical all-zero vector and was assigned the same constant prediction, GO:0004713. Accuracy on this uncovered group was 0.299, simply the proportion of those proteins whose true label happened to be GO:0004713. Conversely, across the 1,945 test proteins containing at least one training-vocabulary peptide, overall accuracy was 0.998. Within individual classes, covered proteins were classified at 0.978 to 1.000 accuracy, whereas uncovered proteins scored 0.000 in every class except the default class GO:0004713 (Figure 2; Table 4). The low overall recall of the peptide arm on GO:0002933 (0.057) and GO:0015977 (0.367) directly reflects that 33 of 35 and 230 of 368 test proteins in those classes, respectively, had zero vocabulary coverage.

**Figure 2.**
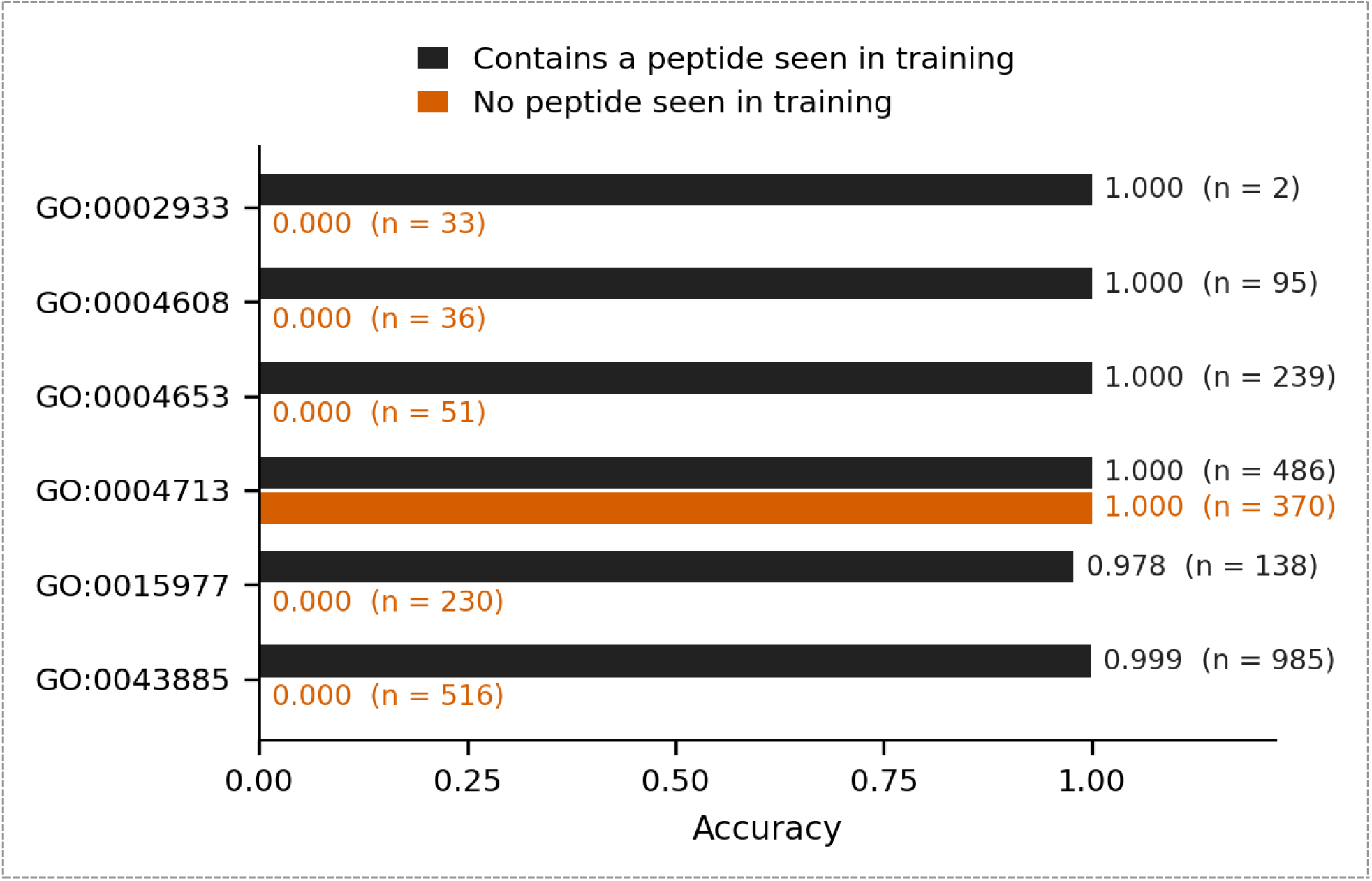
Peptide-arm accuracy depends on vocabulary coverage. Accuracy for each class, separately for test proteins that contain at least one peptide seen in training (black) and those that contain none (orange), with the number of proteins in each group. Proteins with no peptide seen in training all receive the same prediction, GO:0004713, so their accuracy is 1.0 for that class and 0.0 for every other class. Only 2 of the 35 GO:0002933 proteins are covered.

The cause appears to be vocabulary recurrence. Across the corpus, digestion produced 395,350 distinct peptides, of which 81.2% occur exactly once, and the training vocabulary kept 15,304 after the minimum-count threshold. On held-out proteins, 92.2% of peptide tokens were absent from that vocabulary, against 0.003% for the 3-mer arm, whose vocabulary is bounded at 8,792 observed tokens.

Per-class precision and recall reflect the same mechanism (Supplementary Table S3). Peptide precision was at least 0.990 on five of the six classes (and 1.000 on three), with recall ranging from 0.057 to 0.824, whereas GO:0004713 had a precision of 0.497 and a recall of 1.000. Outside the default class, the peptide arm made almost no false-positive calls; when lacking coverage, it defaulted to GO:0004713, where 370 of the 1,236 constant predictions were correct by coincidence (Figure 3A). Accordingly, the peptide arm achieved a higher macro-precision than the 3-mer arm (0.913 vs. 0.848) but a lower macro-recall (0.605 vs. 0.677). In contrast, the 3-mer arm distributed its errors across classes without collapsing onto a default prediction (Figure 3B): excluding the 35-protein class GO:0002933, its precision ranged from 0.428 to 0.966 and its recall from 0.451 to 0.980. Notably, minimal peptide coverage sufficed for near-perfect classification: test proteins with non-zero peptide coverage below 25% were classified at 0.998 accuracy, matching those with coverage above 75%.

**Figure 3.**
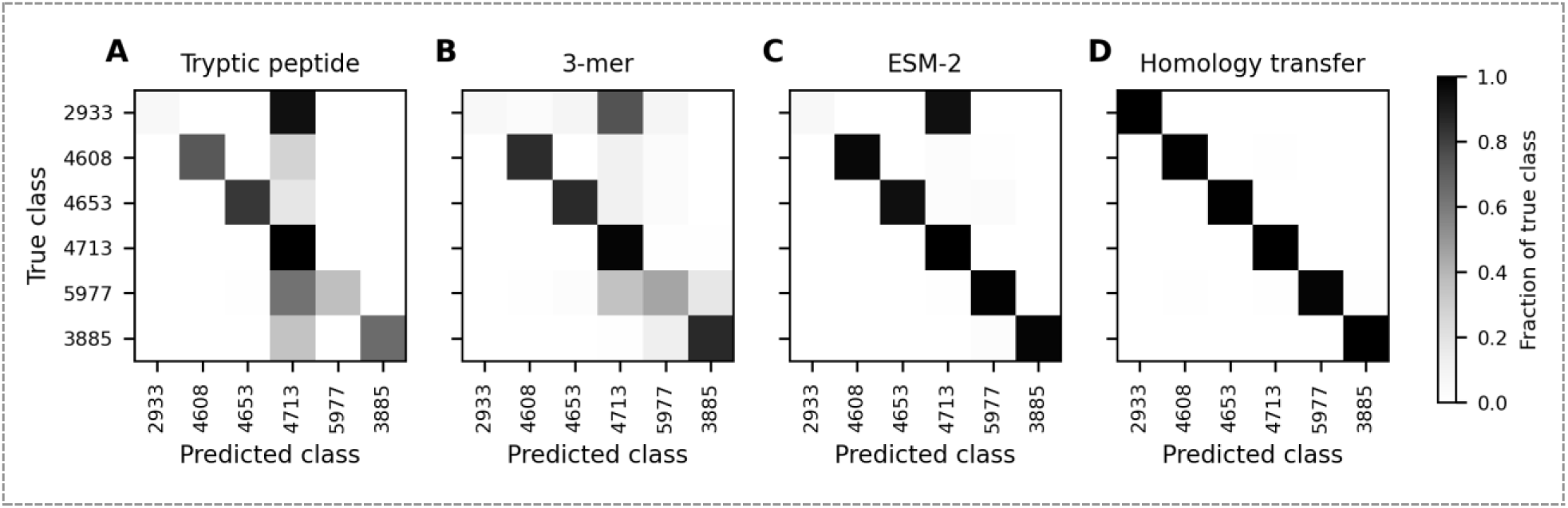
Where each method places its predictions. Confusion matrices on the test set, normalised by true class, for the tryptic peptide arm (A), the 3-mer arm (B), ESM-2 (C) and homology transfer (D). Axis labels give the last four digits of each GO identifier. Peptide errors concentrate in the GO:0004713 column, the default prediction for proteins with no peptide seen in training.

This sparsity is not an artefact of the Word2Vec minimum-count threshold (Table 5; Figure S1). While a threshold of 5 discards 95.4% of training peptides, retaining every singleton (min_count = 1, a 21.7-fold larger training vocabulary of 331,365 peptides) reduced the test token out-of-vocabulary rate only from 92.2% to 83.1%, because peptides observed once in training rarely recur at test time. Lowering the threshold reduced the fraction of uncovered test proteins from 38.9% to 20.8%, but macro-F1 peaked at min_count = 2 (0.665) and was lowest at min_count = 1 (0.559), consistent with prior observations that oversized vocabularies degrade generalization (16).

Overall, peptide-arm accuracy is governed by vocabulary coverage: covered proteins are classified nearly perfectly via an operation resembling exact fragment matching, whereas uncovered proteins receive a uniform guess. Unconditioned aggregate metrics therefore measure vocabulary overlap rather than learned functional generalization.

### 3.3 Peptide topics largely recover protein families

We constructed peptide topics to test whether clustering related peptides into coarser groups could restore recurrence and capture shared functional motifs. If topics represent cross-family functional units, topic clusters should span multiple classes or improve generalization over exact peptides.

We clustered the 15,304 training-vocabulary peptides into 124 topics and assigned each peptide the GO class of its parent training protein to measure topic purity. To identify what drives the most frequent peptides, we ranked peptides by protein occurrence and audited the proportion of fragmentary records among their source proteins.

Rather than capturing cross-cutting functional motifs, topics largely recovered individual protein families and database sampling biases. Of the 124 topics, 95.2% consisted of more than 90% a single GO class, with a median purity of 1.000 (minimum 0.290). Frequency rankings revealed the driver of this concentration: 13 of the top 20 and 31 of the top 100 most frequent peptides derived from a single enzyme; ribulose-bisphosphate carboxylase large chain (rbcL), and topics 27 and 6 consisted predominantly of fragments of that protein. Across the top 200 most frequent peptides, 30.1% of source proteins were fragmentary records, double the 15.0% baseline across the corpus.

Recording out-of-vocabulary peptides in a dedicated 125th count column ensured that no test protein received a zero vector, yet topic performance remained within the peptide arm’s range: macro-F1 was 0.574 (vs. 0.640 for exact peptides), balanced accuracy was 0.601 (vs. 0.605), and macro-AUPR was 0.653 (vs. 0.650). Thus, eliminating zero-vector padding did not close the gap to 3-mers. The topic arm was, however, the only learned representation to predict the smallest class, GO:0002933, at above-chance levels (recall 0.343).

High class purity is partly a consequence of the embedding objective: because skip-gram Word2Vec embeds tokens by within-protein co-occurrence, peptides originating from the same protein family sit near one another in vector space and cluster together. Furthermore, the two-fold fragment enrichment shows that high peptide recurrence reflects sequencing density such as partial rbcL sequences from plant barcoding surveys, rather than conserved functional importance.

### 3.4 Cluster-disjoint splitting leaves substantial identity between partitions

Our evaluation split placed no MMseqs2 cluster in both training and test partitions. To verify whether cluster-disjointness successfully bounded pairwise similarity between partitions, we computed the maximum fractional sequence identity of every test protein to any training protein across alignments covering at least 80% of both sequences.

Cluster-disjoint partitioning did not prevent substantial cross-partition sequence identity. Across the 3,181 test proteins, the median maximum identity to a training protein was 0.450 (interquartile range 0.277–0.651). In total, 70.6% of test proteins shared at least 30% identity with a training protein, 58.9% shared at least 40%, 43.7% shared at least 50%, and 4.5% shared at least 90%. We therefore do not describe this partition as homology-disjoint, and we audit classification accuracy against measured sequence identity below.

### 3.5 Identity effects are confounded by class composition and, for peptides, by coverage

Because sequence distance from the training set is a primary determinant of prediction difficulty (6), classification accuracy would be expected to decline as maximum identity to the training set decreases. We stratified test proteins into bins by their maximum identity to any training protein (including 691 test proteins with no alignment meeting the 80% coverage criterion) and examined accuracy both in aggregate and within individual classes (retaining class-by-identity bins with *n* ≥ 30).

In aggregate, differences across identity bins were heavily confounded by class composition. The bins represent distinct class mixtures rather than comparable populations: the ≤ 30% identity bin is 83.3% GO:0004713 and contains only two classes with at least 10 proteins, whereas the 50–90% identity bin is 70.4% GO:0043885. Consequently, even a trivial baseline that always predicts the single overall majority class varies from 0.008 to 0.215 accuracy across bins.

Evaluating within individual classes removes this compositional confound and shows that accuracy varies only modestly with sequence identity. For the 3-mer arm, accuracy increased with training identity in three of the four classes having multiple bins of *n* ≥ 30 (rising from 0.972 with no alignment to 1.000 at 50–90% identity in GO:0004713; from 0.721 to 1.000 in GO:0004653; and from 0.728 at 30–50% identity to 1.000 above 90% in GO:0043885), but did not increase monotonically in GO:0015977. By contrast, the peptide arm scored 1.000 in every bin of GO:0004713, including the 356 proteins with no 80%-coverage alignment, because GO:0004713 is its default prediction for uncovered proteins.

Conversely, the peptide arm’s sharpest drop to 0.141 accuracy in the 30–50% identity bin of GO:0043885 coincided with 451 of those 526 proteins (85.7%) having no training-vocabulary peptide, whereas ESM-2 achieved 0.998 accuracy on the same bin. Thus, within-class peptide performance is driven by local vocabulary coverage rather than sequence identity.

### 3.6 The findings hold across classifiers

If vocabulary coverage, a property of the input, explains the difference between the tryptic peptide and 3-mer arms, the comparison should hold when the random forest is replaced by other classifiers. We refitted all four representations with logistic regression (standardised features, balanced class weights) and gradient-boosted trees (200 iterations, balanced sample weights) on the same frozen split, without tuning, and compared macro-F1, balanced accuracy, macro-AUPR and ROC-AUC with the random forest (Table 6; Supplementary Table S1).

**Table 6.** Macro-F1 and balanced accuracy of each representation under three classifiers.

| **Representation** | **RF macro-F1** | **RF bal. acc.** | **LR macro-F1** | **LR bal. acc.** | **GB macro-F1** | **GB bal. acc.** |
|---|---|---|---|---|---|---|
| Amino-acid composition | 0.561 | 0.587 | 0.526 | 0.551 | 0.640 | 0.662 |
| 3-mer | 0.690 | 0.677 | 0.683 | 0.697 | 0.746 | 0.739 |
| Tryptic peptide | 0.640 | 0.605 | 0.560 | 0.567 | 0.640 | 0.605 |
| Peptide topics | 0.574 | 0.601 | 0.536 | 0.551 | 0.565 | 0.595 |

*RF, random forest; LR, logistic regression; GB, gradient-boosted trees. Macro-AUPR and ROC-AUC are in Supplementary Table S1.*

Both the 3-mer advantage and the bimodal coverage behavior held across all three classifiers. Overlapping 3-mers outperformed tryptic peptides on macro-F1 under random forest (0.690 vs. 0.640), logistic regression (0.683 vs. 0.560), and gradient boosting (0.746 vs. 0.640), as well as on balanced accuracy and macro-AUPR across every model. ROC-AUC also favored 3-mers under logistic regression (0.945 vs. 0.884) and gradient boosting (0.976 vs. 0.910), while remaining marginally higher for peptides under random forest (0.909 vs. 0.905). Under all three classifiers, the 1,236 test proteins lacking training-vocabulary peptides received a single constant prediction that was correct for 29.9% of the group (Supplementary Table S2), whereas accuracy on the 1,945 covered proteins reached 0.998 with random forest and gradient boosting and 0.927 with logistic regression. The relative ranking of the remaining arms varied with the classifier: under gradient boosting, simple amino-acid composition matched tryptic peptides on macro-F1 (0.640) and surpassed them on balanced accuracy (0.662 vs. 0.605), macro-AUPR (0.722 vs. 0.650), and ROC-AUC (0.949 vs. 0.910).

These results confirm that the coverage bottleneck is a property of the tokenization rather than the classifier.

## 4. Discussion

Our results demonstrate that a sequence tokenization transfers to unseen proteins only when its vocabulary recurs across proteins, and that biological grounding does not by itself guarantee recurrence. We tested the intuitive hypothesis that fragments grounded in biochemistry would generalize better than arbitrary fixed-length windows: tryptic peptides are the physical products of proteolysis (9,10) and the analytes measured in bottom-up mass spectrometry (11,12), whereas a three-residue window is merely an encoding convenience.

### 4.1 Recurrence, not grounding, explains the difference

The 3-mer vocabulary is bounded (8,792 distinct tokens observed), leaving virtually no test 3-mers unseen (0.003%). By contrast, the tryptic peptide vocabulary is effectively open-ended: 23,723 proteins yielded 395,350 distinct peptides, 81.2% of which occurred only once. Because a single amino-acid substitution inside a fragment or at a cleavage boundary creates a distinct token, an exact tryptic peptide acts more like a sequence fingerprint of its parent protein than a reusable building block shared across a family. Consequently, 92.2% of peptide tokens in held-out proteins were absent from the training vocabulary. This bottleneck is not an artefact of frequency pruning or classifier selection: retaining all training singletons lowered the test token out-of-vocabulary rate only to 83.1%, and the 3-mer advantage held across all three classifiers (Section 3.6). These findings align with results from subword tokenization, where algorithms such as byte-pair encoding segment rare sequences into frequent sub-units specifically to guarantee recurrence (13,14,19,20), and where compact vocabularies consistently outperform large ones (16). Enzymatic tokenization fixes token boundaries by cleavage chemistry and leaves token frequency unconstrained. Biochemical grounding and statistical recurrence are therefore distinct properties, and for generalization to unseen proteins, recurrence is the decisive one.

### 4.2 What peptide-based scores represent

Because most tryptic peptides do not recur, peptide-based predictions bifurcate into two extremes. The 61.1% of test proteins containing at least one training-vocabulary peptide were classified at 0.978 to 1.000 accuracy per class under the random forest (0.927 overall with logistic regression), whereas the remaining 38.9% received an identical all-zero vector and were assigned a single default class. Aggregate accuracy on an enzymatic representation therefore measures the proportion of test sequences covered by the training vocabulary rather than the quality of functional inference. Crucially, this vulnerability can be diagnosed before any classifier is trained by computing the fraction of held-out tokens and held-out sequences lacking training-vocabulary coverage; whenever sequence-level non-coverage is substantial, evaluation metrics should be reported separately for covered and uncovered populations.

Moreover, achieving near-perfect accuracy from a small number of shared peptides resembles exact fragment matching—an operation that sequence alignment already performs across the entire test set without coverage failures (homology transfer macro-F1 of 0.995 vs. 0.640 for tryptic peptides). Both homology transfer and ESM-2 (0.997 macro-AUPR, albeit an optimistic bound given pretraining on UniRef (28)) confirm that functional signal is readily recoverable across this split. Pooling related peptides into topics ensured that no protein received a zero vector, yet topic scores remained in the peptide arm’s range because clusters recovered individual protein families and database sampling biases such as abundant *rbcL* fragments, rather than cross-family functional motifs.

These results clarify what is needed to anchor sequence-based function prediction to mass-spectrometry observables. The obstacle is not that tryptic peptides are biologically uninformative, but that exact peptide identity is rarely conserved across distant homologs. A representation that retains enzymatic cleavage boundaries while projecting unseen peptides onto biochemically similar in-vocabulary fragments or decomposing them into shared sub-units could preserve the experimental link without suffering from exact-match sparsity. Evolution-aware subword merges (15) and reduced-alphabet encodings (20) point in this direction, and evaluating such schemes under homology-aware splits is a compelling next step.

### 4.3 Splitting by cluster does not control identity between partitions

Our audit also highlights a broader methodological caveat for benchmark construction. Partitioning sequences by greedy identity clusters is widely used to prevent data leakage between training and test sets (22–24). However, greedy clustering guarantees only that each sequence lies within the threshold distance of its own cluster representative (31,32), not that members of different clusters are distant from one another, because sequence similarity is not transitive. In our corpus, even after programmatically verifying that no 30%-identity cluster crossed the partition boundary, 70.6% of test proteins still shared at least 30% sequence identity with a training protein over an 80%-coverage alignment. Conversely, enforcing strict transitive separation via connected components of the homology graph proved incompatible with class balance, as three of the six GO classes had 42% to 88% of their sequences concentrated within a single component. We recommend that benchmark studies explicitly report the empirical distribution of cross-partition sequence identity—which requires only a single search of test sequences against the training set—and document any trade-off between transitive homology separation and class stratification rather than resolving it silently.

### 4.4 Limitations

Several limitations bound the scope of our conclusions. First, regarding corpus and label structure, our evaluation spans six generally well-separated GO terms including one biological-process term (GO:0015977) that encompasses multiple unrelated folds and biologically overlaps GO:0043885 with test set sizes ranging from 35 to 1,501 proteins. The smallest class (GO:0002933, n = 35*n* = 35was recovered at identical low recall (0.057) across three distinct representations, illustrating a small-sample threshold effect that depresses macro-F1 globally, and 98.7% of the corpus consists of unreviewed TrEMBL records whose annotations are predominantly electronically inferred (2). Second, regarding tokenization parameters, we evaluated a single protease (trypsin) without missed cleavages; while alternative proteases or missed-cleavage tolerances would shift fragment length distributions, the singleton sparsity problem is expected to persist whenever tokens are long enough to be nearly unique (6–40 residues vs. 3-mers). Finally, ESM-2 sequences were truncated at 1,022 residues (primarily affecting GO:0004713), classifiers were evaluated using default hyperparameters without tuning, and cross-identity performance comparisons remain descriptive due to unequal class composition across identity bins.

## 5. Conclusion

Vocabulary recurrence matters more than biological grounding for whether a sequence tokenization transfers to unseen proteins. Although tryptic peptides correspond directly to the chemical products of proteolysis and the experimental observables of bottom-up proteomics, they consistently trailed overlapping 3-mers in Gene Ontology term prediction across all three classifiers tested. Because a single residue substitution alters peptide identity, most tryptic peptides in held-out proteins were never observed during training, reducing classification to near-perfect exact matching on covered proteins and uniform default guessing on uncovered ones. We recommend reporting token- and protein-level vocabulary coverage prior to model training so that vocabulary overlap is not conflated with learned functional generalization. Similarly, because standard cluster-disjoint partitioning left most test proteins with at least 30% sequence identity to a training relative, we recommend auditing and reporting measured cross-partition identity alongside benchmark scores. Bridging experimental proteomics and sequence-based function prediction will require representations that combine enzymatic cleavage boundaries with sub-peptide or substitution-tolerant recurrence.

## Data availability

Code, the full pipeline notebook and all derived result tables are available and archived at [Zenodo DOI: DOI: 10.5281/zenodo.24291728]. Sequence data are not redistributed and can be retrieved from UniProt release 2026_02 using the queries in Section 2.1.

---

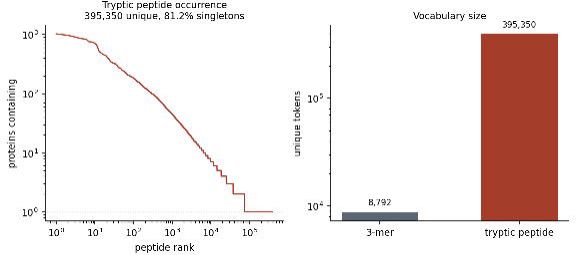

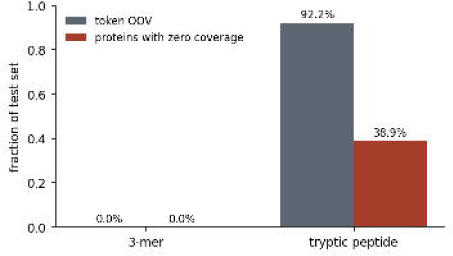

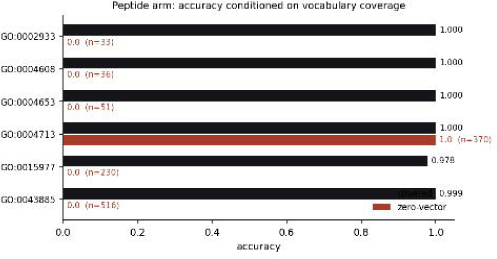

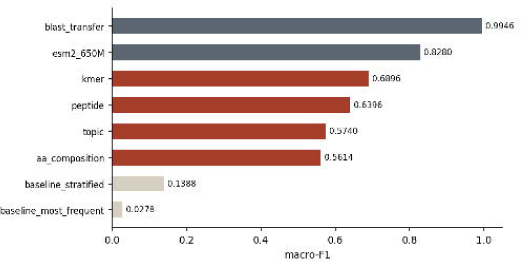

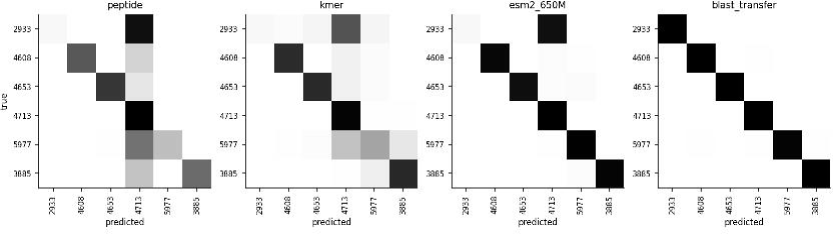

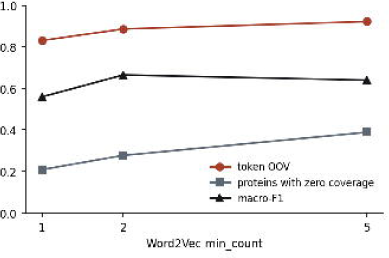

