## Supplementary Table S1 for "Vocabulary recurrence, rather than biological grounding, explains tokenization transfer in protein function prediction"

Tolulope S. Adeyina et al.

### Supplementary Table S1

**Table S1.** Performance of each representation under three classifiers on the frozen test set (n = 3,181). All classifiers used the same split and balanced class weights, with default settings and no tuning.

| **Classifier** | **Representation** | **Macro-F1** | **Balanced acc.** | **Macro-AUPR** | **ROC-AUC** |
| --- | --- | --- | --- | --- | --- |
| Random forest | Amino-acid composition | 0.5614 | 0.5870 | 0.6494 | 0.8764 |
| Random forest | 3-mer | 0.6896 | 0.6771 | 0.7135 | 0.9054 |
| Random forest | Tryptic peptide | 0.6396 | 0.6048 | 0.6501 | 0.9087 |
| Random forest | Peptide topics | 0.5740 | 0.6011 | 0.6526 | 0.8660 |
| Logistic regression | Amino-acid composition | 0.5256 | 0.5506 | 0.6127 | 0.8860 |
| Logistic regression | 3-mer | 0.6825 | 0.6967 | 0.7365 | 0.9452 |
| Logistic regression | Tryptic peptide | 0.5600 | 0.5665 | 0.6186 | 0.8839 |
| Logistic regression | Peptide topics | 0.5363 | 0.5514 | 0.6299 | 0.8597 |
| Gradient-boosted trees | Amino-acid composition | 0.6401 | 0.6620 | 0.7215 | 0.9494 |
| Gradient-boosted trees | 3-mer | 0.7464 | 0.7394 | 0.8006 | 0.9763 |
| Gradient-boosted trees | Tryptic peptide | 0.6402 | 0.6048 | 0.6504 | 0.9101 |
| Gradient-boosted trees | Peptide topics | 0.5648 | 0.5953 | 0.6735 | 0.9150 |

### Supplementary Table S2

**Table S2.** Behaviour of the tryptic peptide arm on test proteins with no peptide seen in training, by classifier.

| **Classifier** | **Test proteins with no peptide seen in training** | **Distinct predictions among them** | **Accuracy, no peptide seen** | **Accuracy, peptide seen** |
| --- | --- | --- | --- | --- |
| Random forest | 1,236 | 1 | 0.299 | 0.998 |
| Logistic regression | 1,236 | 1 | 0.299 | 0.927 |
| Gradient-boosted trees | 1,236 | 1 | 0.299 | 0.998 |

### Supplementary Table S3

**Table S3.** Per-class precision, recall and F1 on the frozen test set, by representation (random forest classifier).

| **Representation** | **Class** | **Precision** | **Recall** | **F1** | **Support** |
| --- | --- | --- | --- | --- | --- |
| Tryptic peptide | GO:0002933 | 1.000 | 0.057 | 0.108 | 35 |
| Tryptic peptide | GO:0004608 | 0.990 | 0.725 | 0.837 | 131 |
| Tryptic peptide | GO:0004653 | 0.992 | 0.824 | 0.900 | 290 |
| Tryptic peptide | GO:0004713 | 0.497 | 1.000 | 0.664 | 856 |
| Tryptic peptide | GO:0015977 | 1.000 | 0.367 | 0.537 | 368 |
| Tryptic peptide | GO:0043885 | 1.000 | 0.656 | 0.792 | 1,501 |
| 3-mer | GO:0002933 | 1.000 | 0.057 | 0.108 | 35 |
| 3-mer | GO:0004608 | 0.966 | 0.855 | 0.907 | 131 |
| 3-mer | GO:0004653 | 0.947 | 0.859 | 0.900 | 290 |
| 3-mer | GO:0004713 | 0.798 | 0.980 | 0.880 | 856 |
| 3-mer | GO:0015977 | 0.428 | 0.451 | 0.439 | 368 |
| 3-mer | GO:0043885 | 0.949 | 0.861 | 0.903 | 1,501 |
| Peptide topics | GO:0002933 | 0.353 | 0.343 | 0.348 | 35 |
| Peptide topics | GO:0004608 | 0.477 | 0.809 | 0.601 | 131 |
| Peptide topics | GO:0004653 | 0.963 | 0.803 | 0.876 | 290 |
| Peptide topics | GO:0004713 | 0.501 | 0.835 | 0.627 | 856 |
| Peptide topics | GO:0015977 | 0.491 | 0.212 | 0.296 | 368 |
| Peptide topics | GO:0043885 | 0.825 | 0.604 | 0.697 | 1,501 |
| Amino-acid composition | GO:0002933 | 1.000 | 0.057 | 0.108 | 35 |
| Amino-acid composition | GO:0004608 | 0.851 | 0.870 | 0.860 | 131 |
| Amino-acid composition | GO:0004653 | 0.765 | 0.776 | 0.770 | 290 |
| Amino-acid composition | GO:0004713 | 0.808 | 0.942 | 0.870 | 856 |
| Amino-acid composition | GO:0015977 | 0.163 | 0.524 | 0.248 | 368 |
| Amino-acid composition | GO:0043885 | 0.936 | 0.352 | 0.512 | 1,501 |

### Supplementary Figure S1

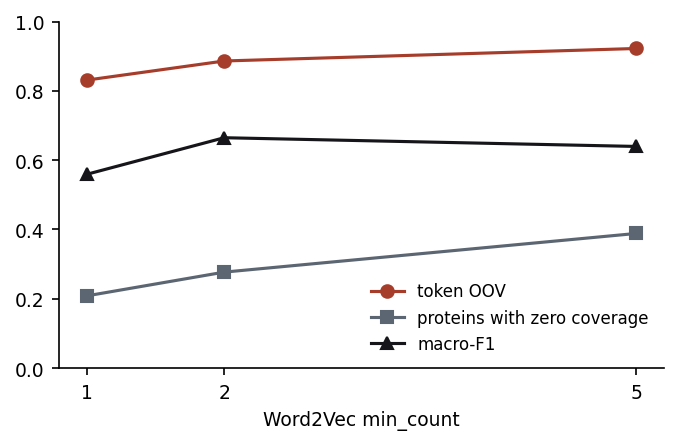

**Figure S1. Effect of the Word2Vec minimum count.** (A) Test tokens unseen in training and test proteins with no peptide seen in training at minimum counts of 1, 2 and 5. (B) Macro-F1 of the peptide arm at the same settings.
